# Automated classification of bat echolocation call recordings with artificial intelligence

**DOI:** 10.1101/2021.06.23.449619

**Authors:** Michael A. Tabak, Kevin L. Murray, John A. Lombardi, Kimberly J. Bay

## Abstract

Acoustic recorders are commonly used to remotely monitor and collect data on bats (Order Chiroptera). These efforts result in many acoustic recordings that must be classified by a bat biologist with expertise in call classification in order to obtain useful information. The rarity of this expertise and time constraints have prompted efforts to automatically classify bat species in acoustic recordings using a variety of learning methods. There are several software programs available for this purpose, but they are imperfect and the United States Fish and Wildlife Service often recommends that a qualified acoustic analyst review bat call identifications even if using these software programs. We sought to build a model to classify bat species using modern computer vision techniques. We used images of bat echolocation calls (i.e., plots of the pulses) to train deep learning computer vision models that automatically classify bat calls to species. Our model classifies 10 species, five of which are protected under the Endangered Species Act. We evaluated our models using standard model validation procedures, but we also performed two out-of-distribution tests. For the out-of-distribution tests, an entire dataset was withheld from the procedure before splitting the data into training and validation sets. We found that our validation accuracy (93%) and out-of-distribution accuracy (90%) were higher than when we used Kaleidoscope Pro and BCID software (65% and 61% accuracy, respectively) to evaluate the same calls. Our results suggest that our approach is effective at classifying bat species from acoustic recordings, and our trained model will be incorporated into new bat call identification software: WEST-EchoVision.

## 1. Introduction

Non-invasive bat survey techniques like passive acoustic sampling have become more commonplace in the past decade, especially with the introduction and spread of white-nose syndrome (Hoyt et al., 2021) across North America and concerns about the possible transmission and spread of COVID-19 into North American bat populations (Platto et al., 2021). Using acoustic recordings to identify bat species has great potential to facilitate research, as many bat species have distinctive call types (Schnitzler et al., 2003). Automated acoustic identification programs offer an objective, repeatable, and rapid means of analyzing large acoustic datasets (Armitage and Ober, 2010; Chen et al., 2020; Hayes et al., 2019; López-Baucells et al., 2019; Mac Aodha et al., 2018). However, these programs have several drawbacks including inadequate bat call identification and reliability (Clement et al., 2014; Gibb et al., 2019; Lemen et al., 2015; Rydell et al., 2017). Specifically, automated bat call classifiers tend to produce significant numbers of false positive identifications (Rydell et al., 2017) and exhibit a surprisingly low level of consistency among programs (Lemen et al., 2015). As a result, most experienced researchers, including the United States (U.S.) Fish and Wildlife Service (USFWS), recommend an experienced acoustic analyst review and validate automated acoustic identification results (Rydell et al., 2017). This recommendation complicates research efforts, as experienced analysts are not available in every research group and their time can be expensive.

We present a novel approach for using deep learning (Goodfellow et al., 2016) to automatically classify bat species in acoustic data. Our method uses plots of frequency over time from zero-cross (ZC) acoustic data to train convolutional neural networks (CNNs) that can automatically classify bat species in acoustic recordings. Our plotting approach isolates pulses from ZC data and creates plots that are classified by computer vision models. This approach provides computer vision algorithms with images of plots similar to those used by experienced bat biologists to manually classify acoustic recordings.

## 2. Methods

### 2.1. Call Library

We used ZC acoustic data to build our call library. ZC acoustic data are low resolution, compared to full spectrum acoustic data, and do not include information on sound amplitude or call harmonics (Parsons and Szewczak, 2009). However, ZC data files are much smaller, use less computer storage space, and can be transferred and analyzed more quickly than full spectrum data files (Britzke et al., 2011). In addition, most bat call classifiers in North America rely on ZC data and the only classifiers certified by the USFWS are those that rely on ZC data. Finally, we had access to ZC data from across the U.S. and to multiple known call libraries of ZC data.

We characterized the bat calls of individual species with two types of bat calls, known calls and expert-identified calls. Known calls were bat calls recorded from bats of known species identity that were visually identified in free flight, light-tagged, hand-released, or recorded near roosts used by a single species (Britzke et al., 2011; Murray et al., 2009). Expert-identified calls originated as passively recorded bat calls that were reviewed and qualitatively identified by an experienced biologist. Qualitative identification entails visual comparison of echolocation call metrics (e.g., minimum frequency, slope, duration) to reference calls of known bats and is a common practice in bat acoustic identification studies (Britzke et al., 2013; Murray et al., 1999; O’Farrell et al., 1999; Russo and Voigt, 2016; Rydell et al., 2017; Yates and Muzika, 2006). The bat call library consisted of call files identified as big brown bat (*Eptesicus fuscus*; EPFU), silver-haired bat (*Lasionycteris noctivagans*; LANO), eastern red bat (*Lasiurus borealis*; LABO), hoary bat (*Lasiurus cinereus*; LACI), gray bat (*Myotis grisescens*; MYGR), little brown bat (*Myotis lucifugus*; MYLU), Indiana bat (*Myotis sodalis*; MYSO), northern long-eared bat (*Myotis septentrionalis*; MYSE), eastern small-footed bat (*Myotis leibii*; MYLE), evening bat (*Nycticeius humeralis*; NYHU), and tri-colored bat (*Perimyotis subflavus*; PESU). In addition, the call library included bat call files that could not be assigned a species identify (“other”) and files that were identified as noise (i.e., non-bat recordings).

### 2.2. Call plots from acoustic recordings

For each bat call, we created an image by plotting the frequency of the call over time (e.g., Fig. 1). ZC files were loaded into R (R Core Team, 2021) using the bioacoustics package v. 0.2.4 (Marchal et al., 2021), which provides, among other information, a vector of acoustic frequencies. We first filtered the data by removing data points with frequencies greater than 125 megahertz (MHz) and less than 16 MHz. Then we identified pulses in the frequency vector, by finding sequences of at least 7 consecutive frequencies of decreasing value. Once the frequency stopped decreasing, it was determined as the termination of the pulse. Using this method to identify pulses also helped to remove noise from the dataset. For each pulse, we calculated the time and frequency of the Unit Invariant Knee (UIK), defined as the point of maximum curvature (Christopoulos, 2016), and colored the UIK orange in the call plots (other points were colored black). We calculated the UIK time and frequency using the R package inflection v.1.3.5 (Christopoulos, 2019). Some acoustic recordings contained many (> 50) pulses, while others only contained few. If a recording contained fewer than five pulses, it was deemed a low quality call, and was not included in model training. We set a maximum of 15 pulses per plot, and if a call contained > 15 pulses, multiple plots for this call were created.

**Figure 1:**
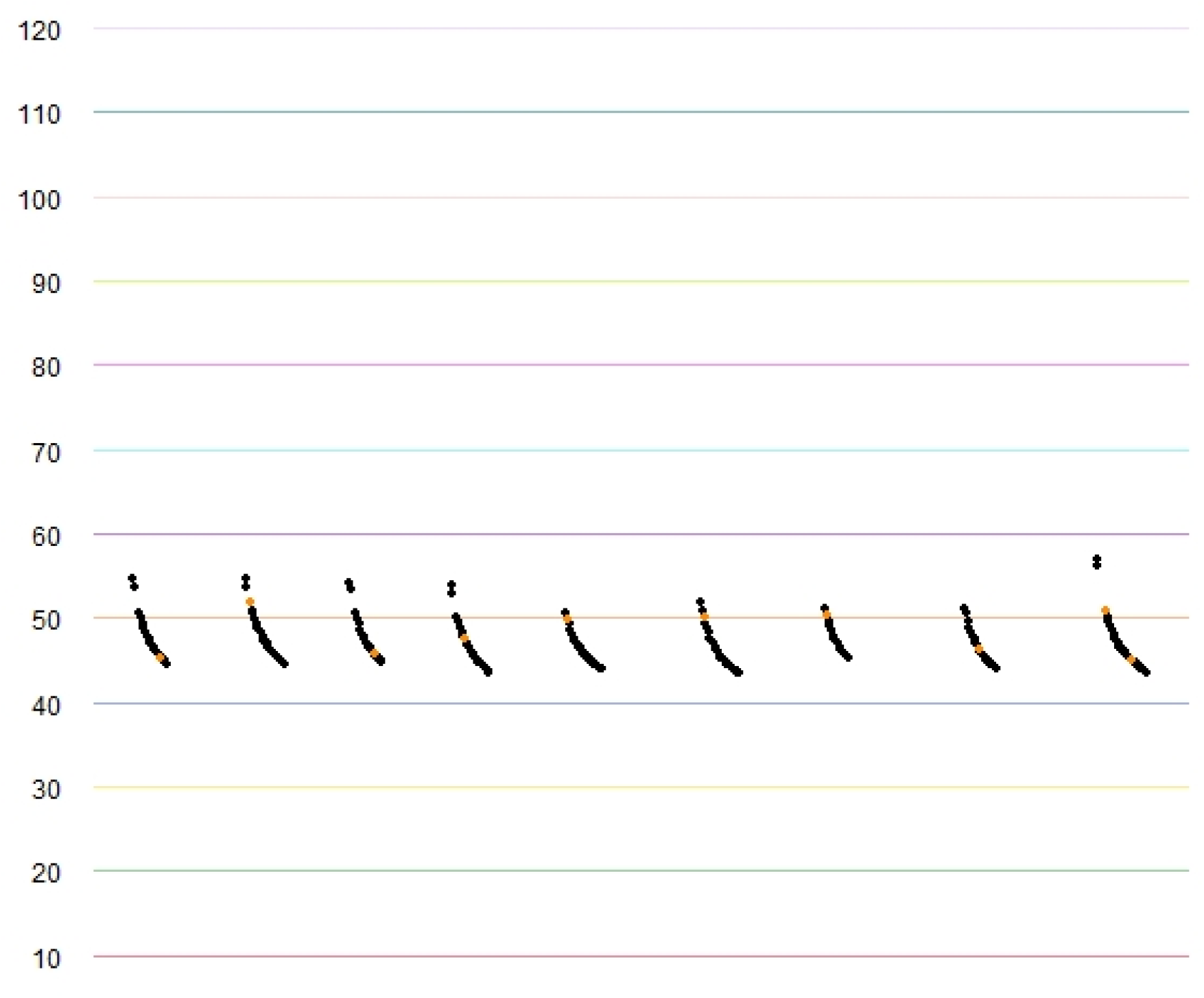
Example of a call plot created to train convolutional neural networks to classify bat calls. This call plot is from an acoustic recording of a tri-colored bat.

### 2.3. Training, validating, and testing the model

To avoid overfitting the model, we excluded one full dataset and used it for out-of-distribution testing. We conducted this procedure two times, where for each experiment we withheld one dataset (Indiana 1 and Illinois 1; Table 1), split the remaining data into training and validation datasets, trained the model, validated the model, and then tested the model on the out-of-distribution dataset.

**Table 1:** Number of bat calls for each species from each dataset. Species codes are presented in the Methods section and in Table 2.

| Project Name | Detector Locations | EPFU | LABO | LACI | LANO | MYGR | MYLE | MYLU | MYSE | MYSO | NYHU | PESU |
| --- | --- | --- | --- | --- | --- | --- | --- | --- | --- | --- | --- | --- |
| Iowa 1 | 2 | 2 | 7 | 15 | 0 | 0 | 0 | 0 | 0 | 0 | 0 | 0 |
| Iowa 2 | 2 | 6 | 11 | 33 | 7 | 0 | 0 | 0 | 0 | 0 | 0 | 0 |
| Iowa 3 | 1 | 1 | 2 | 17 | 3 | 0 | 0 | 33 | 0 | 0 | 0 | 0 |
| Iowa DNR 1 | 4 | 77 | 317 | 86 | 14 | 0 | 0 | 82 | 1 | 0 | 2 | 159 |
| Iowa DNR 2 | 160 | 64 | 514 | 0 | 0 | 0 | 0 | 655 | 818 | 45 | 21 | 116 |
| Iowa DNR 3 | 46 | 51 | 96 | 0 | 0 | 0 | 0 | 114 | 19 | 30 | 3 | 70 |
| Iowa 4 | 2 | 1 | 45 | 4 | 2 | 0 | 0 | 20 | 0 | 1 | 0 | 0 |
| Iowa 5 | 5 | 25 | 47 | 72 | 303 | 0 | 0 | 25 | 2 | 0 | 4 | 0 |
| Illinois 1 | 7 | 136 | 339 | 195 | 258 | 0 | 0 | 115 | 3 | 19 | 4 | 91 |
| Illinois 2 | 8 | 228 | 255 | 47 | 7 | 0 | 0 | 347 | 91 | 290 | 6 | 49 |
| Indiana 1 | 8 | 54 | 80 | 83 | 4 | 0 | 0 | 39 | 11 | 31 | 0 | 250 |
| Oklahoma 1 | 56 | 32 | 1,213 | 24 | 0 | 0 | 0 | 0 | 133 | 0 | 310 | 1,301 |
| Western US Call Library<br>(AZ, CA, NV, SD, UT, WA, WY) | 30 | 29 | 0 | 59 | 70 | 0 | 0 | 40 | 0 | 0 | 0 | 0 |
| Eastern US Call Library<br>(AR, IN, KS, KY, MI, MO, NC, SC, TN, TX) | 77 | 289 | 59 | 101 | 92 | 55 | 6 | 120 | 54 | 164 | 92 | 197 |
| Total (All Libraries) | 408 | 995 | 2,985 | 736 | 760 | 55 | 6 | 1,590 | 1,132 | 580 | 442 | 2,233 |

We used the pulse plots to train CNNs to recognize the bat species in the call. After each testing dataset was withheld, we trained the model on 70% of the classified bat calls, and used the remaining 30% for model validation. The training and validation datasets were selected by stratification of the call label using the Python package SciKit-Learn v.0.23.2 (Pedregosa et al., 2011). After training, we evaluated the model on the validation and testing datasets. Out-of-distribution testing is useful to demonstrate the efficacy of our methods, but by withholding data from training and validation, we reduced our sample size, which is small for some species of conservation concern (e.g., gray bat). Therefore, our final model, which is available in our software (WEST-EchoVision) and is presented in model validation results, does not exclude an out-of-distribution testing dataset when training; it has only training and validation datasets.

### 2.4. Training deep learning models

All CNNs were trained using the ResNet-18 architecture (He et al., 2016), which was obtained from the torchvision package v. 0.8.1 (Pytorch Core Team, 2021). Networks were trained in Python v.3.8.5 (Python Software Foundation, 2020) using Pytorch v. 1.7.0 (Paszke et al., 2019) on an NVIDIA GeForce RTX 3090 graphics processing unit (GPU). We used randomly initialized weights for ResNet in the first epoch. We used a softmax function on the final layer to generate probabilities for each class and selected the class with the highest probability as the classification for each pulse plot. To compare the model prediction to the ground truth classification, we used a cross entropy loss function where each class was weighted inversely proportional to its frequency in the training dataset (species for which more calls were available had lower weights). We used an adaptive movement estimation (Adam) optimization algorithm to update the weights of the CNN at each step. We used a cosine annealing learning rate scheduler, which causes the learning rate to vary over epochs, with an initial learning rate of 10^−4^ and a minimum learning rate of 10^−6^. We trained the models for 15 epochs. In each epoch, if the validation loss was an improvement over the best loss value, the weights from that epoch were saved and loaded in the next epoch.

### 2.5. Model Evaluation

While models were trained and validated at the level of the pulse plot, models were evaluated at the level of the acoustic recording call. (As stated above, multiple plots were occasionally generated from one acoustic recording.) In the rare event plots for the same recording resulted in different predictions (< 5% of all recordings), we took the sum of probabilities of each class for each of the plots from that recording and selected the class with the highest sum of probabilities.

We conducted model validation by using the trained model to evaluate the validation dataset and predict a species for each acoustic recording. We evaluated model performance using several metrics for each class including recall and precision, which summarize the rates of true positives (TP), false negatives (FN), and false positives (FP) and are defined as:

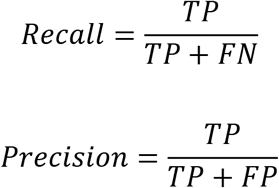

Recall is the proportion of calls of a class that were classified as that class by the model, and precision is the proportion of calls classified as a class that were actually members of that class. We also calculated the *F*_1_ score, which is a summary of these two values:

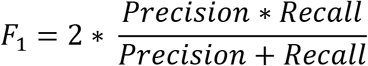

And the accuracy across the whole dataset, which is the proportion of the dataset that was correctly classified.

As we conducted out-of-distribution model testing two times, performance metrics are reported as weighted means. We also evaluated each acoustic recording in these out-of-distribution datasets using Kaleidoscope Pro and BCID. Since these testing datasets were not included in training or validation, they present an effective means for comparing model performance among different software.

## 3. Results

### 3.1. Call Library

We compiled acoustic files from 14 datasets including two existing known call libraries and 12 projects with expert-identified bat calls (Table 1). Bat calls came from 20 states, 122 counties, and 347 unique sampling locations (Appendix). The overall dataset included 17,384 files with 11,514 bat calls files with species identifications and 5,871 files identified as unknown bats or noise. While there were bat calls from 11 species in our call library (Table 1), we only included 10 in model training, as the six calls available for eastern small-footed bats were deemed insufficient for training and validation. With the exception of eastern small-footed bats (six calls) and gray bats (55 calls), the call library included a minimum of several hundred calls from each species (Table 1).

### 3.2. Model Evaluation

Model validation accuracy was 92%. Recall, precision, and *F*_1_ scores were higher for classes for which more training data were available (Table 2). Median testing accuracy for the out-of-distribution datasets in our model was 90%, while Kaleidoscope and BCID software produced accuracies of 65% and 61%, respectively, for the same validation dataset used to evaluate our models (Table 3). Our models outperformed Kaleidoscope and BCID software (as per *F*_1_ score) for all species in the dataset. All of our samples for gray bat were from one dataset (Eastern US Call Library; Table 1), so we could not perform out-of-distribution testing for this species (thus it does not appear in Table 3). Nevertheless, our validation *F*_1_ score for this species was 0.91 (Table 2).

**Table 2:** Model performance on the validation dataset

| **Species** | **Precision** | **Recall** | **F1** |
| --- | --- | --- | --- |
| big brown bat (EPFU) | 0.97 | 0.97 | 0.97 |
| eastern red bat (LABO) | 0.93 | 0.89 | 0.91 |
| hoary bat (LACI) | 0.99 | 0.97 | 0.98 |
| silver-haired bat (LANO) | 0.95 | 0.97 | 0.96 |
| grey bat (MYGR) | 0.95 | 0.88 | 0.91 |
| little brown bat (MYLU) | 0.87 | 0.93 | 0.90 |
| northern long-eared bat (MYSE) | 0.94 | 0.97 | 0.95 |
| Indiana bat (MYSO) | 0.82 | 0.82 | 0.82 |
| evening bat (NYHU) | 0.76 | 0.78 | 0.77 |
| tri-colored bat (PESU) | 0.98 | 0.96 | 0.97 |
| accuracy | 0.92 | 0.92 | 0.92 |

**Table 3:** Comparing out-of-distribution testing performance from our model with those of Kaleidoscope and BCID. The weighted mean scores from the two testing datasets are presented for each species. The first row for a species is from our model, the next row shows the results from Kaleidoscope for that species, and the following row shows the results from BCID. Note that grey bat was excluded from out-of-distribution testing because it was only available in one dataset.

| **Species** | **Precision** | **Recall** | **F1** |
| --- | --- | --- | --- |
| big brown bat | 0.77 | 0.99 | 0.86 |
| (Kaledoscope results) | 0.26 | 0.53 | 0.35 |
| (BCID results) | 0.67 | 0.68 | 0.52 |
| eastern red bat | 0.97 | 0.92 | 0.94 |
|  | 0.88 | 0.32 | 0.36 |
|  | 0.89 | 0.34 | 0.46 |
| hoary bat | 0.97 | 0.95 | 0.96 |
|  | 0.82 | 0.93 | 0.87 |
|  | 0.87 | 0.54 | 0.67 |
| silver-haired bat | 0.99 | 0.73 | 0.84 |
|  | 0.38 | 0.75 | 0.50 |
|  | 0.07 | 0.50 | 0.12 |
| little brown bat | 0.94 | 0.84 | 0.88 |
|  | 0.47 | 0.60 | 0.52 |
|  | 0.60 | 0.41 | 0.46 |
| northern long-eared bat | 0.95 | 0.94 | 0.94 |
|  | 0.21 | 0.24 | 0.20 |
|  | 0.33 | 0.53 | 0.41 |
| Indiana bat | 0.49 | 0.89 | 0.63 |
|  | 0.53 | 0.43 | 0.45 |
|  | 0.43 | 0.79 | 0.54 |
| evening bat | 0.20 | 0.29 | 0.24 |
|  | 0.00 | 0.00 | 0.00 |
|  | 0.00 | 0.00 | 0.00 |
| tri-colored bat | 0.96 | 0.99 | 0.97 |
|  | 0.96 | 0.88 | 0.92 |
|  | 0.87 | 0.80 | 0.83 |
| accuracy | 0.90 | 0.90 | 0.90 |
|  | 0.65 | 0.65 | 0.65 |
|  | 0.61 | 0.61 | 0.61 |

Confusion matrices (Figs. 2–3) from each training and validation exercise demonstrate how each call from each species was classified. The x-axis contains the label that was provided by a human expert (“ground truth”), and the y-axis contains the model prediction for that call. The diagonal of the matrix contains the number of correct predictions for each class. A perfect model would only have numbers on the diagonal. The most common misclassifications, as depicted in these confusion matrices were among eastern red bats and little brown bats, and little brown bats and Indiana bats.

**Figure 2:**
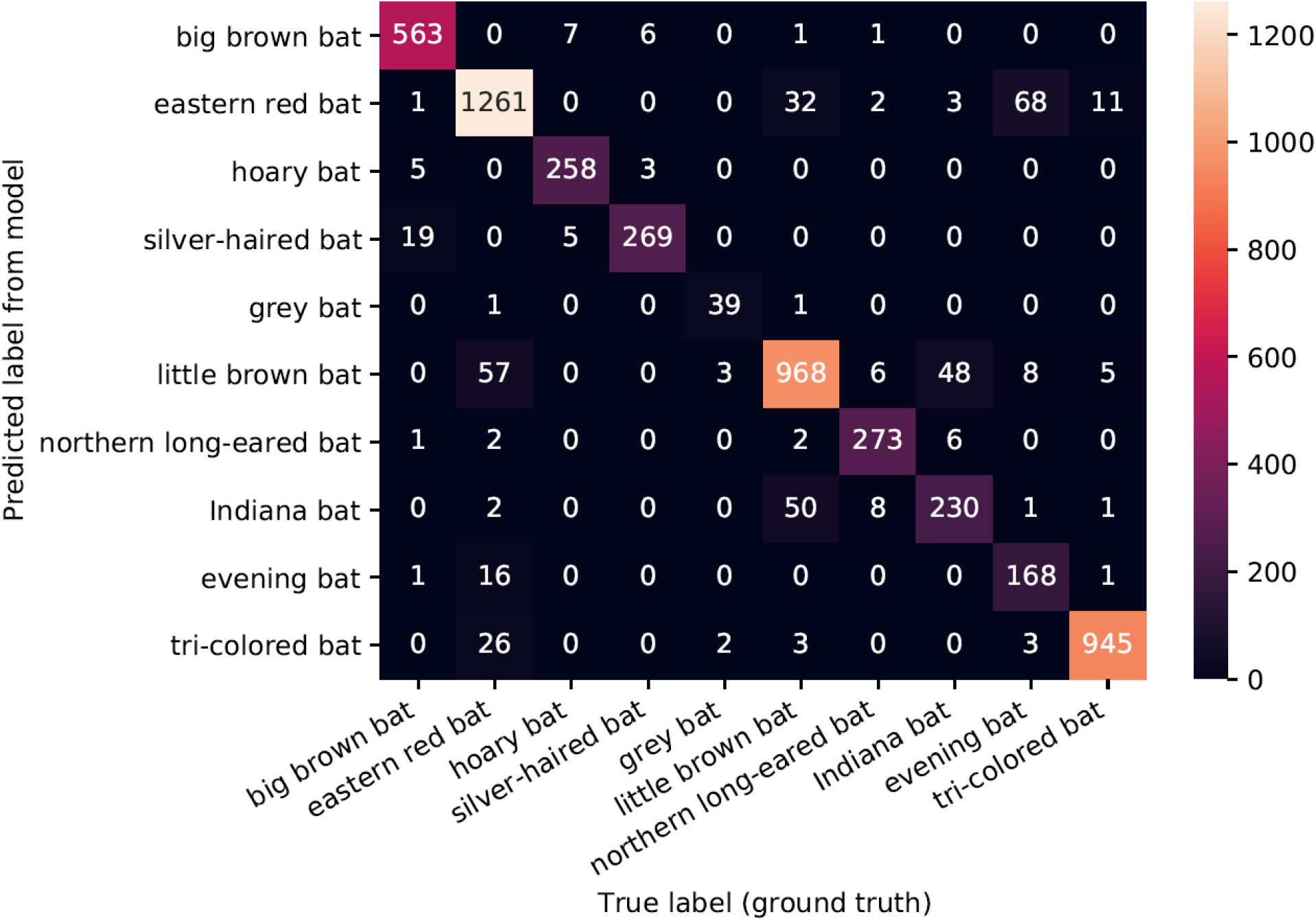
Confusion matrix for the model when the Indiana 1 dataset was excluded from training and used for testing.

**Figure 3:**
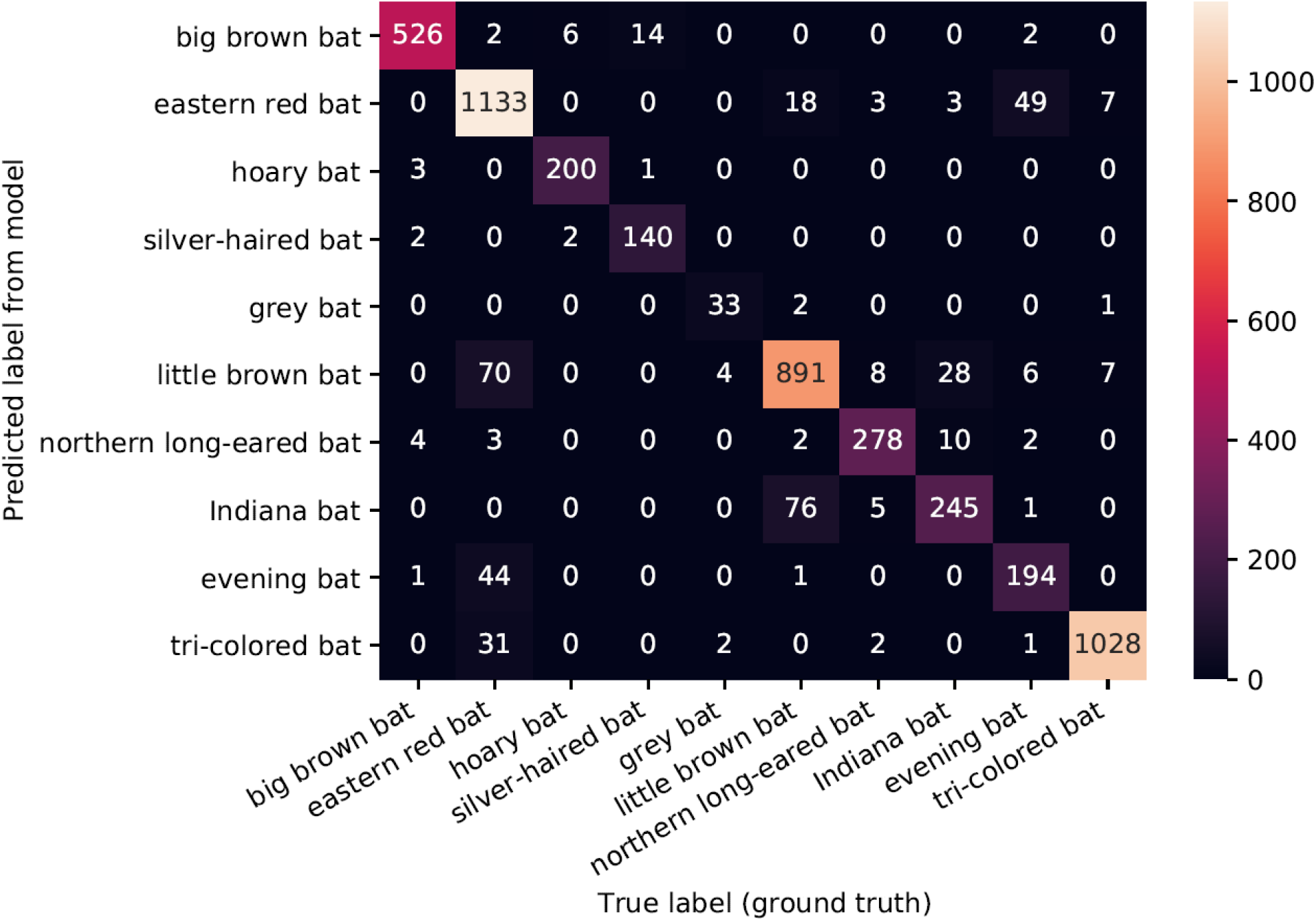
Confusion matrix for the model when the Illinois 1 dataset was excluded from training and used for testing.

## 4. Discussion

We found that our models classified bat species with relatively high validation accuracy as well as out-of-distribution testing accuracy. This indicates that our approach is effective for classifying bat species from their acoustic recordings and that it will generalize to other datasets. Our models predicted species in acoustic recordings with higher accuracy than currently available software, even when we compared our results on out-of-distribution testing data to results from other software. For species of conservation concern, it is especially important to be able to detect them accurately. In our dataset there are five species of conservation concern: Indiana bat, grey bat, and little brown bat are listed as endangered (Arroyo-Cabrales and Ospina-Garces, 2016; Solari, 2018a, 2017; United States Fish and Wildlife Service, 2021a, 2021b), northern long-eared bat is threatened to extinction (Solari, 2018b; United States Fish and Wildlife Service, 2021c) and tricolored bat is vulnerable to extinction (Solari, 2018c). Validation and out-of-sample accuracies for these species in our models outperformed those of Kaleidoscope and BCID software (Tables 2 and 3).

Our methods can be used to automatically classify bat species using their acoustic recordings and thus help efforts to conserve bat species of conservation concern. For example, “smart curtailment” algorithms reduce turbine speed at wind farms when collisions with bats are more likely (Hayes et al., 2019). Our software can be used to measure bat activity near wind turbines, providing more information for researchers developing smart curtailment strategies, and thus further reduce bat fatalities at wind farms while maximizing wind energy production. Furthermore, this technology can also be used to classify acoustic recordings in real time at wind farms and advise curtailment when species of conservation concern are active near turbines. Finally, our software can be used to automatically classify acoustic recordings and allow for the expansion of acoustic monitoring programs that aim to study the abundance and distribution of bat species. By improving our understanding of their ecology, we can help preserve species of conservation concern (Hey et al., 2003).

Deep learning is increasingly being used to aid ecological research and obtain information from non-invasive sampling (Ruff et al., 2020; Tabak et al., 2019). Our results suggest that deep learning architectures commonly applied to computer vision tasks can also be used to classify species in acoustic recordings. Often, computer vision species classification models perform poorly when applied to out-of-distribution datasets (Schneider et al., 2020; Tabak et al., 2020). This likely results from problems distinguishing animals from the background of the image (i.e., their surrounding environment), especially in camera trap images (Beery et al., 2019; Singh et al., 2020). Our plotting approach removed some of the background noise, such that the model can train based only on the pulses of the acoustic recording. Thus, the algorithm is analyzing plots in a similar manner to what human experts find useful when classifying bat species from echolocation plots.

While our method can be useful in future efforts to classify bat species from acoustic recordings, there is room for improvement, as our models’ performance on out-of-distribution testing data was imperfect, especially for those species with fewer acoustic recordings available (Tables 2 and 3). One way to improve our model is incorporating more training data for species where we have small sample sizes (e.g., gray bat) or to develop a model with higher resolution, full-spectrum acoustic data. Furthermore, by incorporating more species into our training dataset, we could produce a model that recognizes more bat species and is more generalizable.

## 5. Conclusions

We applied a novel computer vision approach to analyzing bat species in acoustic recordings. Our model is capable of automatically classifying bat species in acoustic recordings. It outperforms commercially available software for all species analyzed in this dataset. This technology can be applied in studies of bat distribution and abundance, and it can also be used to inform smart curtailment strategies aimed at reducing bat collisions with wind turbines. Our software will soon be available in WEST-EchoVision.

## Abbreviations

ZC: Zerocross
CNN: Convolutional Neural Network
USFWS: United States Fish and Wildlife Service

## 6. Acknowledgments

MidAmerican Energy Company funded the development to help support their efforts in developing a Habitat Conservation Plan for wind energy generation assets in Iowa. We thank the Illinois-Iowa Ecological Services Field Office and Iowa Department of Natural Resources for access to acoustic data and the USFWS “Cooperative Endangered Species Conservation Fund” that supported the collection of acoustic data in Iowa. We also thank all of our collaborators for access to acoustic data from multiple states and to those who helped collect and compile known bat calls including Dr. Lynn Robbins, Dr. Eric Britzke, Ryan Allen, and Andrew Krause.

## Notes

### Competing Interest Statement

The authors have declared no competing interest.

